# The lifetime cost of reproductive potential – who spends the most?

**DOI:** 10.1101/2021.03.22.432784

**Authors:** Shai Fuchs, Miki Goldenfeld, Michal Dviri, Clifford Librach, Micha Baum

**Affiliations:** Lunenfeld Tanenbaum Research Institute, Mount Sinai Hospital, Toronto, Ontario, Canada; Faculty of Medicine, Tel Aviv University, Tel Aviv, Israel; CReATe Fertility Centre, 790 Bay Street, Suite 1100, M5G1N8, Toronto, Canada; Department of Obstetrics and Gynecology, University of Toronto, 1 King’s College Circle Toronto, M5S 1A8, Canada; Institute of Medical Sciences, University of Toronto, 1 King’s College Circle Toronto, M5S 1A8, Canada; Department of Physiology, University of Toronto, 1 King’s College Circle Toronto, M5S 1A8, Canada; Chaim Sheba Medical Center, Tel-HaShomer Hospital, Ramat Gan, Israel

## Abstract

**Objectives:** To determine who spends more energy over a lifetime on maintaining their reproductive potential: men or women?

**Design:** As a model and energetic equivalent, we set the mass of gametes supported over time from birth until exhaustion of fertility. We calculated gender-specific dynamics of gamete pool mass over time. To this purpose we collated data from existing literature, accounting for gamete volume over stages of development, time in each stage, mass density, and count. Our model generates the integral, or area under the curve (AUC) of the gamete pool mass over a lifetime as a proxy to energetic requirements.

**Main outcome measures:** The area under gamete mass curve over a lifetime in men and women.

**Results:** The number of gametes over a lifetime is 600,000 in women and close to 1 trillion in men. Accounting for mass and time, women invest approximately 100 gram*days in maintaining the female oocyte pool. Women reach 50% of lifetime AUC by age 10, and 90% by age 25. Men invest approximately 30 Kg*days over a lifetime (300-fold more), reaching 50% of lifetime AUC at age 37 and 90% at age 62 years old.

**Conclusions:** The study quantifies for the first time the area under gamete mass in men and women through a nuanced calculation accounting for all components of post-natal gamete dynamics. We found a 300-fold excess is supported male gamete mass over a lifetime (100g*days vs. 30 Kg*days in females vs. males, respectively). Our methodology offers a framework for assessing other components of the reproductive system in a similar quantitative manner.

## Introduction

Reproduction is a fundamental feature of all living organisms. In humans, as in other organisms, life history can be formalized as strategies to optimize the allocation of resources to reproduction(1). Sexual reproduction comes with an energetic price tag. Trade-offs in energy allocation between growth, reproduction, and survival lead to pivotal tradeoffs for fecundity across the animal kingdom(1),(2).

Reproductive efforts can be grossly divided into two main components: 1) the pre-fertilization phase (gamete production, maintenance and transport, hormonal regulation, menstrual cycle including ovulation, and mating efforts), representing mainly the aspect of energy spent on the reproductive potential, which is not translated to offspring biomass. 2) the post-fertilization phase, or direct investment in the offspring (gestation, childbirth, lactation, and parenting efforts)(1).

The post-fertilization phase vastly dominates the female reproduction-related energetic expenditure(1). The reported long-term effects of reproduction on female longevity and health are consistent with that. One study in preindustrial women population, reported reduced survival and higher susceptibility to infections after the age of 65 in mothers of twins, as compared with mothers delivering singletons. This risk has been further elevated when these women started reproducing early (3). A more recent study supported the observation that in women, higher early-life fecundity (<25 years), was associated with increased overall mortality risk (4). Another study showed that lower parity and older age at last birth are associated with decreased risk of female mortality past the age of 60 when compared to their non-late fertile counterparts. The wives’ longevity was more sensitive to the effects of parity and late fertility than their husbands’ (5). An additional study concluded that women had a significantly higher risk of mortality than men at all levels of parity and worsened with increasing parity (6). A recent study found a trade-off between reproduction and post-reproductive lifespan in females, but not in males, based on a large cohort of more than 118,000 individuals (7).

It is thus the human male vs. female pre-fertilization energetic costs which represents an intriguing research question. This is especially challenging as the reproductive potential is achieved through strategies that radically diverge between the sexes. In females, oogenesis occurs during fetal development, forming a finite number of stored gametes that are used periodically over a defined reproductive lifetime. A pool of 6-7 million oocytes at around five months of gestational age, declines continuously until menopause(8, 9). In contrast, male germ cells migrate to the genital ridge during embryogenesis and undergo a series of mitotic divisions by puberty. Spermatogenesis is initiated at the onset of puberty, supporting the continuous production of approximately 100-200 million spermatocytes every day(10), and men can produce and emit sperm throughout their lifespan(11).

Analysis of vertebrate models suggested that the striking surplus of male gametes translates to excess energy investment by males(12). On the other hand, human data suggest that gametogenesis does not represent a significant reproductive energy requirement(1). This is supported by studies demonstrating that sperm counts, motility, and viability are not associated with moderate energy limiting conditions(13-15).

In this paper, we aim to compare the order of magnitude of energy directly invested in gamete pool generation, maintenance, and renewal between human males and females. We address gender-specific gamete pool mass over time, accounting for conservation and renewal by comparable energetic currency. Our model studies the area under the curve (AUC) of the gamete pool mass over a lifetime as a proxy for energetic requirements. A nuanced quantitative approach based on the existing body of literature is employed to answer our research question – *who spends the most on their respective gamete pool: men or women?* Through eliciting the components of our calculation, we aim to divulge underrecognized insights and highlight existing knowledge gaps. We find whether the gender-specific cost of reproductive potential is unbalanced in a manner that is in keeping, or at odds, with the toll of pregnancies. Our findings could contribute to the modeling of fertility-associated energetic trade-offs in men vs. women. Finally, establishing an order of magnitude ratio between the energy cost of the male vs. female gamete pool may help to elicit differences between the sexes in the susceptibility of their reproductive potential to energy limitations, and to serve further research efforts.

## Methods

We refer to a unit mass supported over unit time (g*day) as an energetic currency equivalent. This simplification allowed us to compare a gradually depleting pool of relatively large oocytes to a constantly replenishing pool of much smaller spermatocytes, and account for the change in mass that occurs in both populations during maturation. We, therefore, aimed to compare the integral over time (or area under the curve) of gamete biomass between sexes from birth.

To that end, we searched the literature for quantitative properties pertaining to oocyte and sperm parameters relevant to our calculation. In females, this included: A. non-growing follicles: count at birth and dynamics of atresia over life. B. Growing follicles: total recruited over a lifetime for each stage and duration of time in that stage, measures of oocyte volume and mass density. In males, this included: A. The spermatogonial stem cell (SSC) pool size at birth and over life. B. During spermatogenesis: clone size, cell size, and duration of time on each stage, allowing us to calculate the mass· time area under the curve (AUC), attributable to a single spermatozoon found in the semen. Then, we coupled that with approaches to reach a nuanced estimate of sperm production over a lifetime. We narrowed our scope to gamete measures only; therefore, our model and calculations do not account for the supporting tissues such as granulosa, theca, and Sertoli cells.

## Results

### Lifetime supported oocyte mass

As a background to our calculation, we note the physiological framework: the female gametes differentiate during the fetal period and are maintained until menopause as a pool of oocytes with the potential to progress to ovulation in response to hormonal cues. Each oocyte is embedded in a follicle - a structure of supporting cells and matrix. At any time point, a single oocyte may either progress in maturation or senesce and undergo atresia(16). The key literature underlying the quantitation of oocytes starts with the pioneering work of Eric Block in the early 1950s (17, 18), and relies on histological studies of ovaries recovered from donors of various ages.

### Dynamics of the Non-growing follicles pool over life

Follicles containing oocytes arrested in the diplotene stage of meiosis I constitute the ovarian follicular reserve, providing lifetime reproductive potential(19). Wallace and Kelsey published in 2010 a meta-analysis aggregating histological samples of 325 ovaries across a wide age range (0.6 to 51.0 years), from which they were able to derive a model that described the age-related decline in non-growing follicle (NGF) number. They report a median reservoir of 300,000 oocytes per ovary(20). This estimate is within the range of 266,000 to 472,000 independently calculated for the mean number of NGF at birth(16). Based on the exponential fit for the decay of the NGF pool formulated by Wallace and Kelsey (Figure 1A), we calculated annual AUC from birth to age 50 of NGF number. Thus, for AUC of the number of oocytes supported each day from age 0 to 50 we calculate:

**Figure 1.**
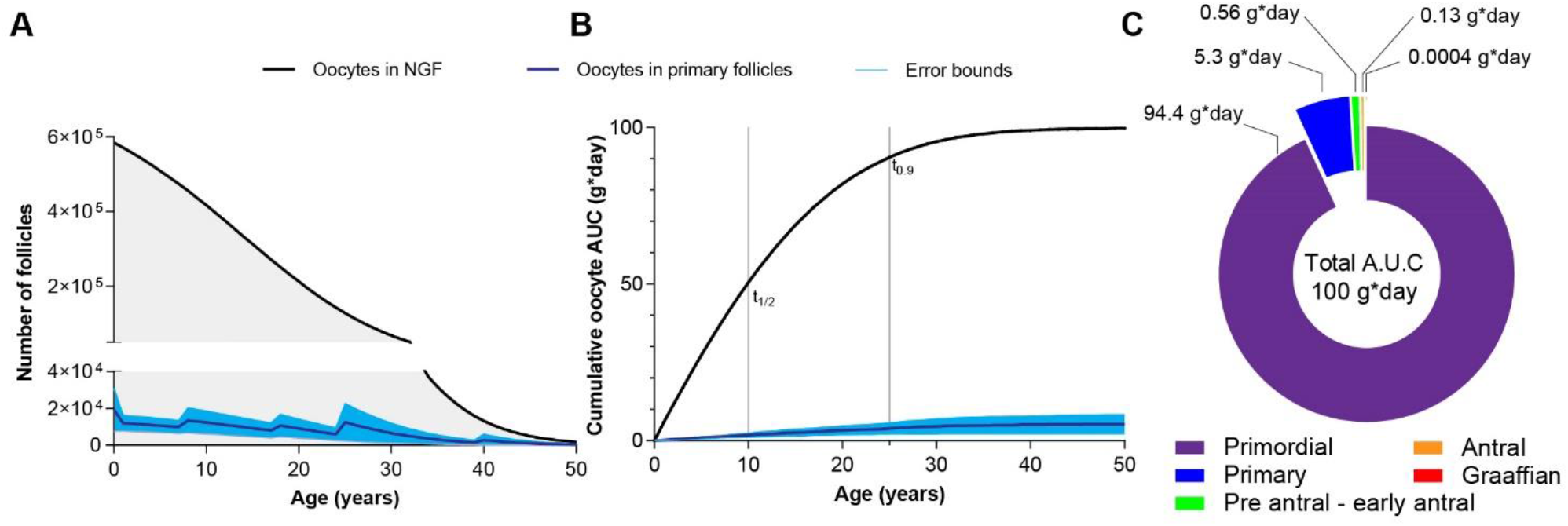
Depleting oocyte pool and cumulative AUC of oocyte mass over a lifetime. A. Decay of non-growing follicle population following Wallace and Kelsey model is used as a foundation for our calculation of oocyte mass supported over time. NGF includes oocytes in primordial and primary follicles, which differ in size. Age-specific mean (purple line) ±SD (blue range) of primary oocyte fraction (Supplementary S1 tab *Fraction of primary follicles*). Based on that an annual supported mass of oocyte in NGF can be calculated (Supplementary S1 tab *Fraction of primary follicles*). Panel B presents the cumulative AUC of the supported oocyte in NGF mass over life, vertical markers represent the age where 50% and 90% of lifetime oocyte AUC are accumulated. Panel C represents total lifetime supported mass*time for oocytes in primordial, primary, and through stages of growing follicles.

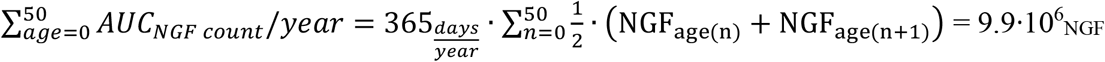

In other words, if we were to count oocytes in non-growing follicles in both ovaries daily from birth to menopause, and sum the counts, we would expect to be at the range of 10 million oocyte days (Supplementary S1 tab *non-growing follicles*).

### Primary Follicle fraction in NGF

By most accounts, the NGF pool is composed of both oocytes in primordial follicles and primary follicles(21, 22). The mean oocyte diameter increases during the transition from an oocyte in a primordial follicle to an oocyte in a primary follicle from 36μm to 42μm, respectively(23). Therefore, the fraction of oocytes in primary follicles needs to be accounted for separately, owing to the different oocyte mass.

We collected studies that altogether analyzed ovaries from 68 women aged 0-50 and reported the fraction of primary follicles(17, 18, 24-26). Aggregating by age groups we derived the mean fraction of primary follicles per age. Using mean+1SD as an upper bound of the estimated fraction of primary follicles, we found that it is below 5% in the neonatal, child, pubertal and young woman (ages 0-24 years), 15% at 25-39 years, and up to 50% at ages 40-49 years (Supplementary S1, tab *Fraction of primary follicles*, Figure 1B). Next, based on a calculated mean volume of 2.44 · 10^−8^*ml* for a primordial oocyte increasing to 3.88 · 10^−8^*ml* for a primary oocyte, and using age-specific primary oocyte fraction, we calculated a weighted average volume of an oocyte in the NGF by age. Multiplying the days of supported oocyte count by the age-appropriate mean oocyte volume, provides the total volume of oocytes in NGF supported each year (Supplementary S1).

Since there are no direct measurements of human oocyte mass or mass density in the literature, we refer to oocyte mass density of 1.05g/ml in porcine oocytes(27) which we used as the best available approximation to human oocytes.

Thus, supported NGF mass over a lifetime was calculated as follows:

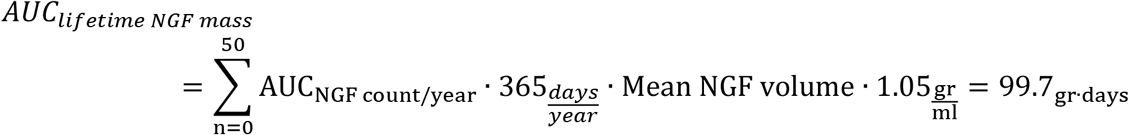

Of note, if we calculate cumulative supported oocyte mass in primary follicles alone, we found a lifetime cumulative mass of 5.3±3.1 _gr· days_ (supplementary S1 tab *Fraction of primary follicles*)., Thus, nearly 95% of the lifetime NGF mass is in the supported primordial follicles (Figure 1B, C).

If we assume energy invested in supporting oocytes is proportional to the area under the curve, the median value is reached at age 10. Furthermore, according to these calculations, by the age of 25, a woman will have invested 90% of her lifetime energy in supporting the NGF pool (figure 1B).

### Maturing follicles

Oocytes depleted from the NGF pool, belong either to follicles that had undergone atresia or to follicles that are progressing to maturation. This multi-stage progression is characterized by changes in the size of the follicle, number, and morphology of cells surrounding the oocyte, and cell cycle stage(21). For our calculation, we accounted for each stage separately, considering the number of oocytes entering it, stage-specific oocyte mass, and time spent per stage (supplementary S1 tab Maturing follicle)(16, 21, 23, 24, 28).

In two detailed accounts of progression of ovarian follicular development in primates, Gougeon et al, were able to quantify the fraction of follicles entering each of the eight classes of maturing oocytes, representing stages of maturation from pre-antral to preovulatory. His analyses provided transit time per stage(21) and the fraction of follicles lost to atresia at each stage(16). Our literature search did not reveal a systematic count of oocytes entering each stage over time.

However, given the rate of loss to atresia at each stage, and given that from class 8, oocytes progress to ovulation, an estimate of the total number of ovulatory cycles over lifetime provided a basis for a retrograde calculation. For example, given that 50% of the oocytes in class 7 progress to ovulation(21), the lifetime number of oocytes in class 7 is twice the number of ovulatory cycles. This calculation can then be extended to calculate the number of oocytes in class 6 (where 77% of the oocytes undergo atresia)(21) and so on.

The number of lifetime ovulatory cycles (LOC) can be crudely estimated to be about 400, based simplistically on the mean number of years from menarche to menopause times 12 cycles/year. A more nuanced approach, taking into account months of oral contraceptive use, pregnancy, breastfeeding, and missed or irregular cycles, would result in an average LOC of 325 ovulatory cycles in a lifetime(29).

Based on 325 oocytes reaching ovulation and the approximate loss to atresia in each cycle leading up to ovulation, we can deduce an estimation for the number of oocytes at each stage of maturation over a lifespan. We accounted for the corresponding mass of oocytes in each stage based on the mean diameter per stage(23). Remarkably, about 20,000 oocytes are recruited in growing follicles throughout life. Due to their fast maturation and high rates of atresia, the total mass of oocytes in growing follicles supported over lifetime amounts to 0.7_g*days_ (Supplementary S1 tab *growing follicles*).

Thus, lifetime total supported oocyte mass is maintained at ∼100_g*days_, of which about 99% goes towards supporting oocytes in NGFs (Figure 1C).

### Lifetime supported sperm mass

What is the male gamete mass supported over a lifetime? A male is born with a cohort of SSC. Each undergoes a series of mitotic divisions estimated to reach over 1 billion SSC at puberty(30). From the onset of spermarche, spermatogenesis, culminating in cohorts of mature spermatozoa, takes place continuously throughout male adult life.

### Lifetime sperm production

Our calculation relies on approximating the number of sperm cells produced over life. This can be approached by knowing the production rate per gram of parenchyma over a lifetime.

Alternatively, the same outcome can be reached by multiplying the number of emissions over a lifetime by the mean sperm count per ejaculation. We employed both approaches independently to validate the estimate reached.

### Number of ejaculations in a lifetime

Spermarche, the first spontaneous emission of sperm was determined by studies based on analysis of urine samples, thus capturing spontaneous emission. Overall, the reported age of onset is at a range of 13.0-14.0 years, with a mean of 13.4 years (31),(32),(33),(34),(35).

What are the dynamics of male ejaculation frequency over life? Very few studies address this question, mainly in association with measures of prostatic health. The single large-scale study on this subject is a cohort study based on recall of ejaculation frequency(36). In this study, a cohort of 29,342 men with a mean age of 58.9±2.5 years reported their ejaculation frequency in the preceding year and was categorized based on frequency category. Participants were also asked to recall ejaculation frequency on their 3rd and 5th decades of life. The average monthly ejaculation frequency recalled was 15.1 for ages 20-29, 11.4 for ages 40-49, and 7.2 per month at age 59, in keeping with a linear decline over advancing age (Supplementary S2 tab Ejaculation frequency). Linear fit for the mean monthly frequency by age yields the following equation for ages 20-59:

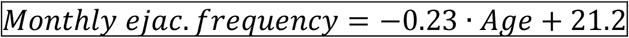

A smaller cohort described by Brody *et al* provides an independent validation to this approximation. The study cohort of 42 healthy, non-obese men aged 20-40 (mean 25.3±4.0), included prospective and recall data on masturbation and intercourse frequency. This cohort reported a mean of 14.9 ejaculations per month(37). This correlates very well with the mean frequency projected from the linear fit that for a 25-year-old man predicts a monthly frequency of 15.4 ejaculations per month(36).

We, therefore, used this linear model spanning ages 14-70, as data suggests a 10-fold drop in sperm count in men 70 and older(11). This resulted in an estimation of 7,800 ejaculations in a male lifetime.

### Sperm count

Since sperm count declines with age, we sought literature that provided age-related sperm count values(11) (38) (39, 40). Collating the data points from these resources, and assuming a linear fit, we reached a projection for age-related sperm count in adults (Supplementary S2, table Sperm count):

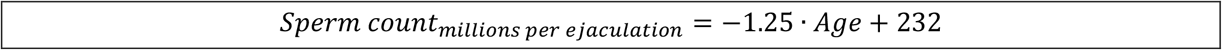

Our implied assumption that the decline of sperm count is linear and monotonous is a simplification(11). However, projections based upon this model receive independent reassurance through the WHO report (2010)(41), a key reference for semen characteristics. It reports a median sperm count across the entire study population (ages 17-67, mean 33±5) to be 196*10^6^ sperms per ejaculation. This median is in keeping with the mean count predicted by the linear fit for age 33, which is 190 million sperm per ejaculation.

Thus, based on the linear fit for ejaculation frequency and sperm count we can calculate (x=age):

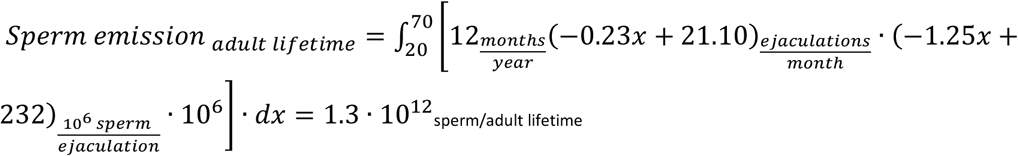

This outcome is comparable with a crude calculation based directly on the WHO mean sperm count per ejaculation:

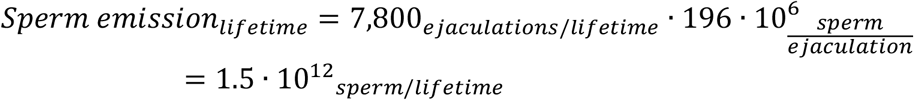

### Sanity check: re-calculating based on measured daily sperm production

The calculation of lifetime sperm output can be complemented by independent literature regarding daily sperm production (DSP). Based on autopsies of 10 men aged 20-29, Johnson et al. reported DSP= 8.5±1.3 · 10^6^ per gr parenchyma, leading to DSP=149±32 · 10^6^ per testis(42). A cohort of 132 men, aged 21-79 reports an age-related decline in sperm production(43). A linear fit for age-related decline in DSP projected it to be *DSP*_*millions per gr parenchyma*_ = −0.065 · *Age* + 7.7

However, the calculated DSP by Johnson relied on a count of the spermatids, divided by the duration of the spermatid stage in spermatogenesis which at the time was considered to be 2.9 days. Based on a more updated review, the round spermatid stage lasts 8 days, which leads to a need to update the cited accounts by 2.9/8=0.3625 fold. Incorporating that correction factor, we calculated:

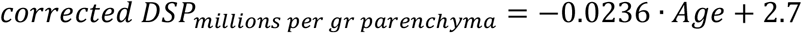

Testicular parenchyma weight did not change with age and was stable at 80% of testicular weight yielding 15.3±0.4g, and 13.8±0.4g in the right and left testicles respectively, or ∼29g parenchyma in total. Thus, for x=age we calculated:

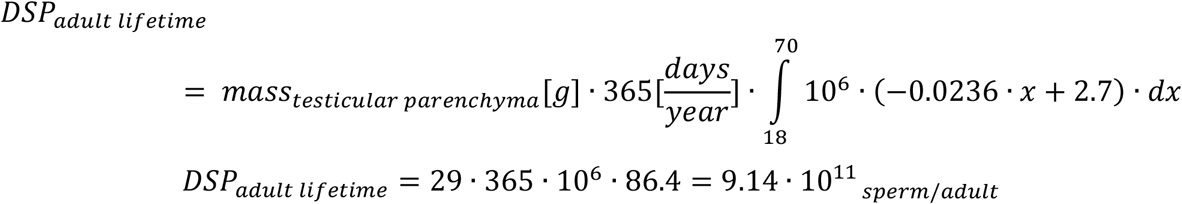

To account for the years of puberty we extrapolated from the rate of growth of testicular volume. A recent cross-sectional study of 514 healthy Norwegian boys provided measures for the linear part of the testicular growth curve based on both orchidometry and sonographic measures(44). Assuming parenchyma fraction and DSP per gram parenchyma to be stable during puberty, the sperm production between ages 14-18 is 7.4· 10^10^, thus contributes little to the lifetime count, which can now be summarized as:

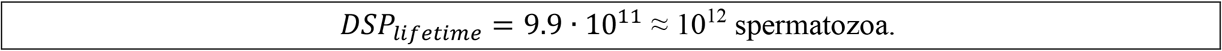

### Sperm cell mass supported over time

Through two independent accounts, we were able to determine that approximately 1 trillion sperm cells are produced over a male lifetime. To translate this into mass supported over time, we determined AUC supported during differentiation of a single spermatozoon.

Spermatogenesis initiates with SCC undergoing mitotic divisions (clonal expansion), which culminates in 4 primary spermatocytes(45). Then, two meiotic divisions take place, as well as the loss of approximately half of the daughter cells to atresia(46), leading to 8 spermatids. Finally, the spermatids differentiate into mature spermatozoa transiting through the epididymis (Figure 2)(45).

**Figure 2.**
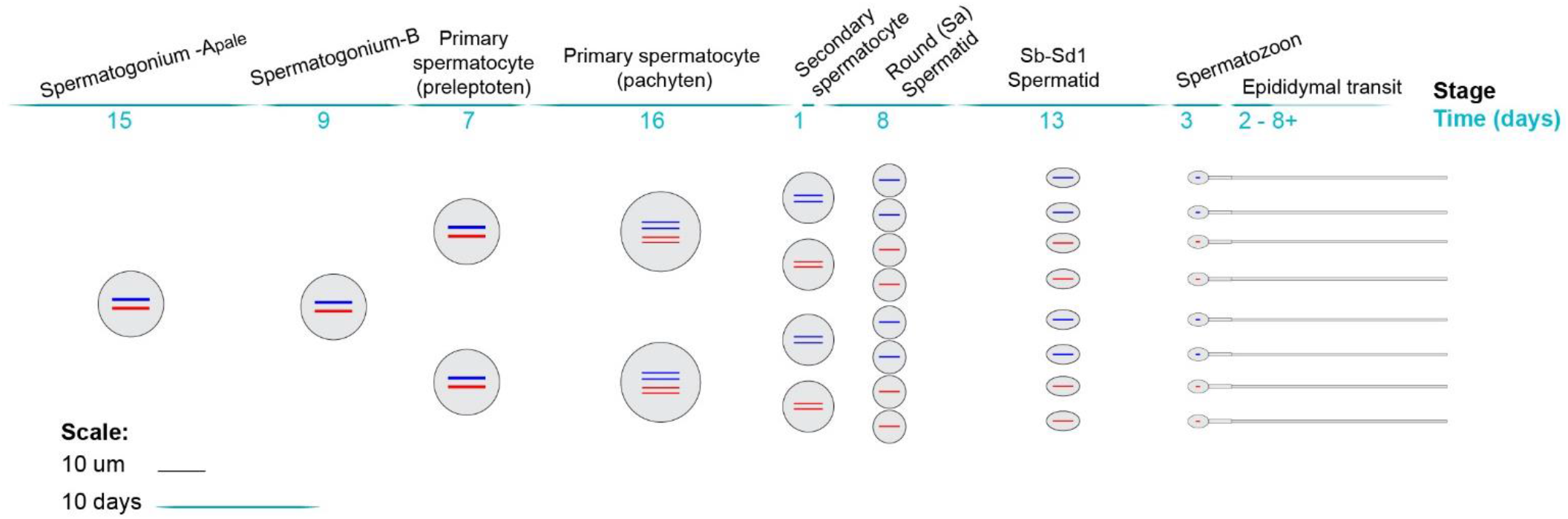
Sperm developmental stages for calculation of supported gamete mass. Visualization of the clonal expansion in number and size from spermatogonium to spermatozoon. The count, time, and size scales correspond to the data reported in table 1 and detailed in supplementary S2 tabs Sperm volume, Sperm developmental stages.

A cohort of spermatogonia commits to spermatogenesis every 16 days, and the total length of the process was initially estimated at an average of 74 days(47). A more recent comprehensive review maintained this mean of 74 days with a 95% CI of 69-80 days^(45)^. To account for the mass supported over time at each stage, per final spermatozoon, we used reported size measures to calculate volume per stage (Supplementary S2 tab *Sperm volume*)(48-50).

For most of the developmental stages described above, the measures available are diameters from which a volume can be derived. However, with few exceptions, a direct measurement of gamete mass is not available. In an account of rabbit and bovine spermatozoa, their mass density was determined to be at a range of 1.1-1.3gr/ml(51). Density in earlier stages of sperm development was not directly assayed. However, based on water content it can be estimated to be closer to the density of most somatic cells at 1.1gr/ml(52). We, therefore, used 1.1gr/ml as an approximation for human sperm mass density during development.

Finally, we divided the mass supported at each stage by a factor representing the expansion in progeny from that stage to spermatozoon. This yielded the mass supported at each stage per one spermatozoon (Supplementary S2 *Sperm developmental stage*). Table 1 summarizes the dimension and count properties for the stages of spermatogenesis.

**Table 1:**
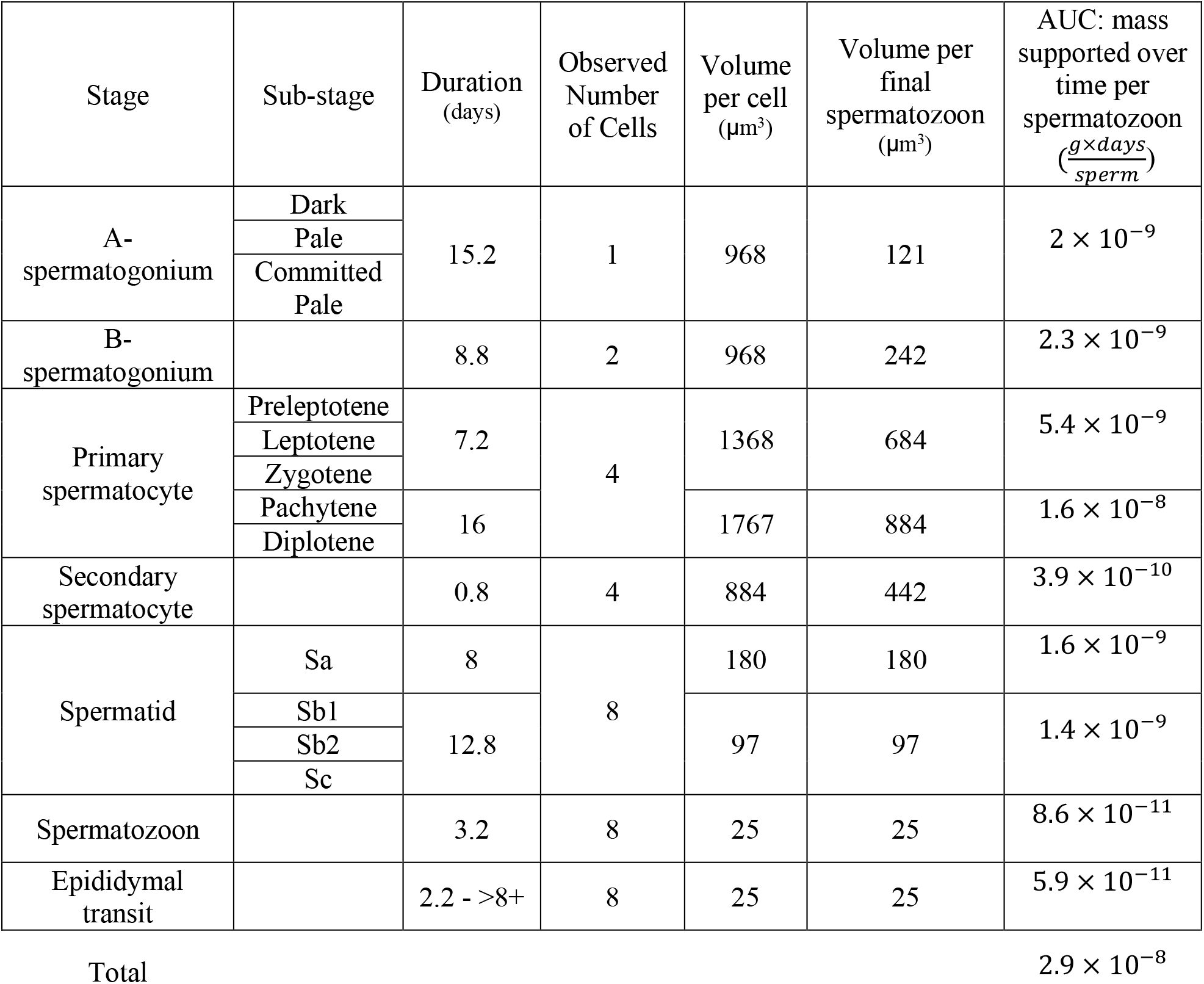
Mass supported over time per final spermatozoon. Each spermatozoon is a part of a clone. Table 1 illustrates the data used to calculate the area under the curve of mass supported over time per final spermatozoon. Cell volume per given stage is based on measured diameters of published representative cells (detailed account in supplementary S2 tab *sperm volume*). At each stage, clone size is affected by division and atresia events. Volume per cell is therefore multiplied by the ratio between clone count at a given stage and the final clone count of spermatozoa. This yields the volume supported per final spermatozoon. Finally, the volume supported per final spermatozoon is multiplied by the approximated mass density of 1.1g/ml and by the duration of that stage to provide area under the curve of the mass supported over stage duration per final spermatozoon. The sum of AUC for all developmental stages projects 2.9· 10^−8^g*days supported per spermatozoon (detailed calculations and references in Supplementary S1 tab *Sperm developmental stages*).

### Spermatogonial stem cells

A male is born with a population of SSC which proliferates through mitotic divisions until puberty. We used a meta-analysis, aggregating data from 334 boys aged 0-14, which provided an age-specific 95% CI for SSC per milliliter of testicular tissue(53). Testicular volume is essentially unchanged in the first five years of life(54). From a cross-sectional study of 452 boys aged 6-16, we extracted an age-specific testicular parenchyma volume during pre-puberty(44). By multiplying the SSC density per unit testicular volume by age-appropriate testicular volume, we delineate the expansion of SSC from birth to over 1 billion cells at the onset of spermarche, which corresponds with previously published estimates(30, 55).

### The area under the curve of supported sperm mass over a lifetime

Based on mass supported over time per sperm cell, and the number of cells produced every year, we generated an annual and lifetime AUC for the supported male gamete mass (Figure 3A, B). When applying lifetime sperm production into this calculation, we preferred to use the outcome which relied on the daily sperm production, since that underlying literature relies on a more robust methodology (histological samples vs. retroactive questioners).

**Figure 3.**
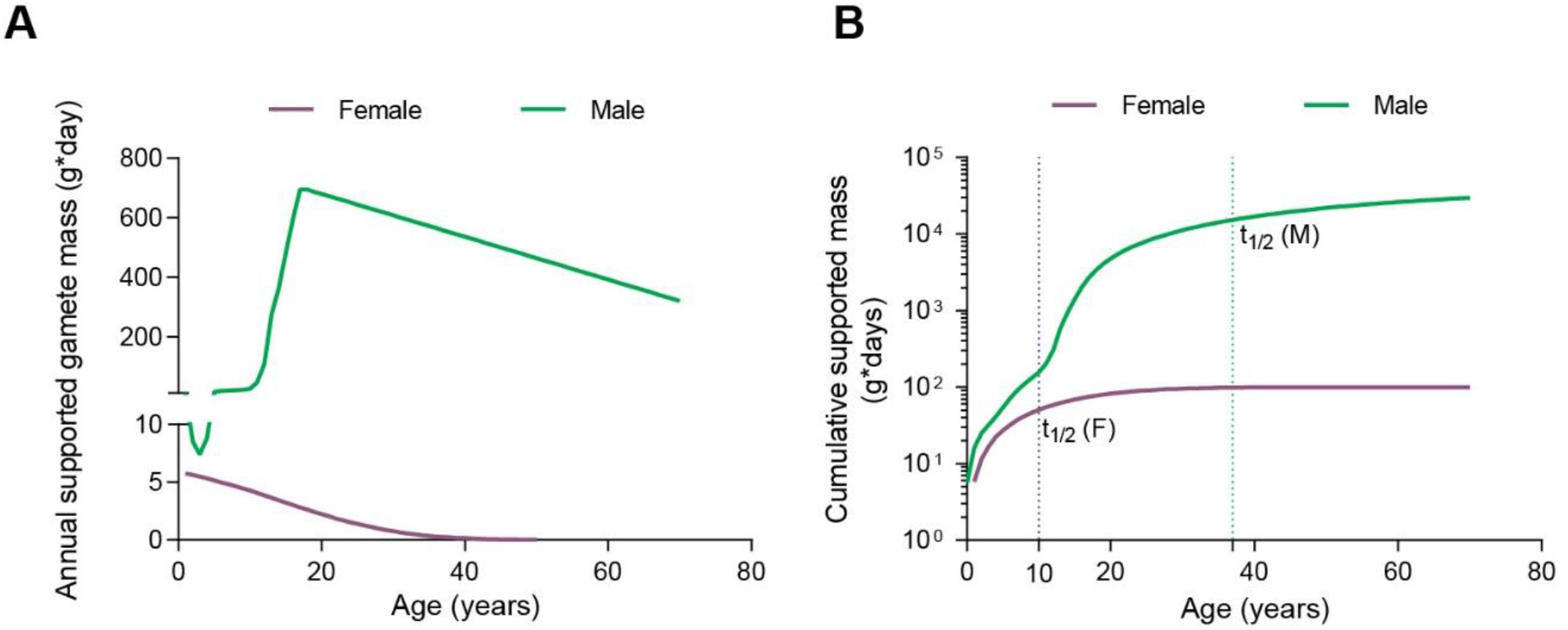
Annual and cumulative gamete mass male vs. female. Annual mass supported over time decays monotonously in females, corresponding to a gamete pool of about 600,000 at birth the undergoes gradual atresia. In males the congenital pool of SSC undergoes expansion to over 1 billion until puberty, then the onset of spermatogenesis leads to a brisk rise in annual gamete mass followed by decay which we approximated to linear (supplementary S2). The cumulative gamete mass AUC (panel B) for males and females over a lifetime, assuming negligible contribution of ages 50+ and 70+ in females and males respectively. Vertical lines denote the age at which 50% of lifetime cumulative gamete AUC is reached in females (10 years old) and males (37 years old).

Accounting for SSC supported mass in prepuberty, and total supported mass during spermatogenesis from age 14 to 70, we found that in a male lifetime, the energetic expenditure to support the gamete mass is approximately 30_Kg*day_ (Supplementary S2 tab *cumulative supported mass for age*). This estimate places the pre-fertilization energetic cost for men at 300-fold higher than the 100_g*day_ equivalent in women. The dynamics of the cumulative energetic investment are also distinct compared to women. These include a spike in supported gamete mass in mid-puberty which is not demonstrable in girls. Men also cross half of their lifetime energetic cost far later than women (at age 37 vs. 10 respectively) (Figure 3).

## Discussion

Maintaining reproductive potential is a fundamental evolutionary task, achieved by strikingly different solutions across various species(1, 2). The energetic cost of this potential in humans represents a fascinating query, yet exceedingly difficult to quantitate.

Human male and female gamete pools diverge by number, size, and temporal dynamics. The asymmetry in gamete counts (male dominance) and size (female dominance) were each analyzed separately, serving opposing claims on the dominant energetic toll of reproductive efforts(12, 56, 57)

Our study is the first to focus on quantifying the overall differences in the dynamics of maintaining the gamete pool in men versus women. To fully account for the costs of male and female reproductive physiology, multiple components should be considered and quantified. These may include costs of hormonal secretion and signaling schemes, maintenance of stable thermal environment for gonads, supporting tissues in the reproductive organs, cyclicity, and even differences in muscle and bone mass. For simplicity, these complex components were left out of our model, a model based solely on daily gamete headcount, accounting for mass.

Is our energetic coin of choice - the AUC of gamete mass, representative of the degree of overall energy invested towards reproduction? Secretion of sex steroids – absolute levels, and dynamics of cyclicity appear to be more strongly correlated with external energy limitation and with fertility outcome(1, 58).

However, a crude measure with some resemblance is effectively used to compare the energetic cost of reproduction between species. The gonado-somatic index was suggested to be proportional to energetic investment in reproduction across six marine fish species(59). This provides support to the notion that within species overall mass of the gamete pool in gonads may also represent the overall energetic investment. Focusing our inquiry into a detailed juxtaposition of a key component of the reproductive system would also help establish principles for assessing other components of that system in a similar quantitative manner. Furthermore, as with any question approached through a more nuanced quantitative approach, the investigation itself exposes knowledge gaps.

Our methodology addresses the most fundamental quantitative properties of a biological system – number count, mass, time. As previously demonstrated in other domains of human physiology, addressing these basic components exposes gaps in existing literature and highlights underrecognized facets of the system(60, 61)

The dynamics of pre-fertilization energy investment are quite striking. We calculated the t_50_ and t_90_, namely the age by which a person will have accumulated 50% and 90% respectively of their lifetime area under the gamete mass curve. Remarkably, in women, these milestones are reached at age 10 years for t_50,_ i.e. well before the age at menarche (median age 12.25 years in the U.S)(62). Female t_90_ is at age 25, thus younger than the mean maternal age at first birth in the United States, currently 26.3 years(63). In men, we note t_50_ and t_90_ to be at ages 37 and 62, respectively. This is although men are born with SSC at a number estimated to be at least 300,000 cells per gonad by conservative estimates(30) and up to millions of SSC. This is a similar range to NGF count at birth in the ovaries(20). It is therefore the proliferative dynamics that determine both the males’ far larger energy investment, as well as far slower dynamics of exhausting their reproductive potential.

Our knowledge on timelines associated with fertility, menopause, and andropause is in keeping with the time scales predicted by the cumulative area under the gamete mass curve. The age-dependent decline in gonadal function is indeed faster in women; whereas, in men, it is far more gradual, and shows a high degree of inter-individual variability(64-66). Moreover, the gamete mass dynamics also correlate with the differences in ages of menopause and andropause. By the age of 54, 85% of women will reach menopause(67), whereas only 30% of men in their 50s will experience symptoms of andropause caused by low testosterone levels(68, 69)

When it comes to the basic physical properties of the developing human gamete, we found the existing literature quite depleted. There is a lack of data on direct mass and mass density measurements for almost all human gamete developmental stages, apart from spermatozoa. In an era where the transcriptional signature of each cell type is available, this serves as a reminder that well-validated quantitative physical properties of key cells still await systematic characterization. Similarly, of the physiological properties we investigated, age-dependent changes in sperm production and emission are only partially characterized(43, 44). Our analysis, based on DSP data, estimates roughly 1 trillion (9.9· 10^11^) sperm cells in a lifetime. Reassuringly, a recently published independent account reached a very similar estimation of 9.25· 10^11^spermatozoa(55). In comparison, our calculation based on ejaculation frequency and mean sperm count yielded an estimate that was 50% higher. This serves as an indication that age-dependency of the frequency of emission as well as mean sperm count needs to be characterized with greater stringency. This aspect is of particular growing interest, in light of accumulating evidence for a decline in sperm count in recent decades in the western hemisphere(70-72),(73) This indicates a need to revise old datasets, and address age, geography, and ethnicity in a more comprehensive manner. Particularly since histologic studies pertaining to daily sperm production, atresia (degeneration) during spermatogenesis, and their change with age rely on a body of literature generated 30-40 years ago(46, 49, 74).

Our model for the energetic cost of the gamete pool findings that the toll on men is at a range of 300-fold higher than that of women. It remains to be seen whether this parameter serves as a proxy to sex differences in overall pre-fertilization reproductive energy effort. However, it does offer a nuanced quantitative approach that can and should be implemented across other domains of reproductive physiology.

## Copyright statement

The Corresponding Authors have the right to grant on behalf of all authors and do grant on behalf of all authors, a worldwide license to the Publishers and its licensees in perpetuity, in all forms, formats, and media (whether known now or created in the future), to i) publish, reproduce, distribute, display and store the Contribution, ii) translate the Contribution into other languages, create adaptations, reprints, include within collections and create summaries, extracts and/or, abstracts of the Contribution, iii) create any other derivative work(s) based on the Contribution, iv) to exploit all subsidiary rights in the Contribution, v) the inclusion of electronic links from the Contribution to third party material where-ever it may be located; and, vi) license any third party to do any or all of the above.

## Supporting information

Supplemental information S1 Oocyte parameters

Supplemental information S2 Sperm parameters

## Acknowledgments

This research was conceived during the course “Cell Biology by the Numbers” by Professor Ron Milo at the Weizmann Institute of Science, Israel. We thank Prof. Milo for illuminating discussions during the early conceptualization of our study.

## Contributors

S.F. and M.G. contributed equally to this paper. S.F and M.G conceptualized the study idea and methodology and constructed the quantitative model. S.F, M.G, and M.D. wrote the first draft of the manuscript. Multiple versions of the manuscript were critically revised by all authors. The analysis and findings were validated by M.B and C.L. All authors have approved the manuscript’s final version to be published.

## Transparency declaration

The lead author* affirms that this manuscript is an honest, accurate, and transparent account of the study being reported; that no important aspects of the study have been omitted; and that any discrepancies from the study as planned (and, if relevant, registered) have been explained.

- *The manuscript’s guarantor.

